# *Mycobacterium tuberculosis*-specific CD4^+^ and CD8^+^ T cells differ in their capacity to recognize infected macrophages

**DOI:** 10.1101/262493

**Authors:** Jason Yang, Daniel Mott, Rujapak Sutiwisesak, Yu Jung Lu, Fiona Raso, Britni Stowell, Greg Babunovic, Jinhee Lee, Steve Carpenter, Sing Sing Way, Sarah Fortune, Sam Behar

## Abstract

Containment of *Mycobacterium tuberculosis* (Mtb) infection requires T cell recognition of infected macrophages. Mtb has evolved to tolerate, evade, and subvert host immunity. Despite a vigorous and sustained CD8**^+^** T cell response during Mtb infection, CD8**^+^** T cells make limited contribution to protection. Here, we ask whether the ability of Mtb-specific T cells to restrict Mtb growth is related to their capacity to recognize Mtb-infected macrophages.

We derived CD8**^+^** T cell lines that recognized the Mtb immunodominant epitope TB10.4_4-11_ and compared them to CD4**^+^** T cell lines that recognized Ag85b240-254 or ESAT6_3-17_. While the CD4**^+^** T cells recognized Mtb-infected macrophages and inhibited Mtb growth in vitro, the TB10.4-specific CD8**^+^** T cells neither recognized Mtb-infected macrophages nor restricted Mtb growth. TB10.4-specific CD8**^+^** T cells recognized macrophages infected with *Listeria monocytogenes* expressing TB10.4. However, over-expression of TB10.4 in Mtb did not confer recognition by TB10.4-specific CD8**^+^** T cells. Importantly, CD8**^+^** T cells recognized macrophages pulsed with irradiated Mtb, indicating that macrophages can efficiently cross-present the TB10.4 protein and raising the possibility that viable bacilli might suppress cross-presentation. Importantly, polyclonal CD8**^+^** T cells specific for Mtb antigens other than TB10.4 recognized Mtb-infected macrophages in a MHC-restricted manner.

As TB10.4 elicits a dominant CD8**^+^** T cell response that poorly recognizes Mtb-infected macrophages, we propose that TB10.4 acts as a decoy antigen. Moreover, it appears that this response overshadows subdominant CD8**^+^** T cell response that can recognize Mtb-infected macrophages. The ability of Mtb to subvert the CD8**^+^** T cell response may explain why CD8**^+^** T cells make a disproportionately small contribution to host defense compared to CD4**^+^** T cells. The selection of Mtb antigens for vaccines has focused on antigens that generate immunodominant responses. We propose that establishing whether vaccine-elicited, Mtb-specific T cells recognize Mtb-infected macrophages could be a useful criterion for preclinical vaccine development.

## Introduction

Unlike most disease-causing pathogens, *Mycobacterium tuberculosis* (Mtb), the cause of tuberculosis (TB), persists in humans because of its highly evolved ability to evade and subvert the host immunity [1]. Mtb subverts vesicular trafficking, prevents phagolysosome fusion, and replicates in an intracellular niche within macrophages, allowing it to evade detection by humoral immunity [2]. Mtb also delays the initiation and recruitment of T cell immunity to the lung, promoting the establishment of a persistent infection [3]. Despite these challenges, T cell immunity does occur and plays an essential role in controlling the infection in both mice and humans [3-5]. With 10 million new TB cases annually, an effective vaccine would offer a cost-effective way to prevent TB and attenuate this persistent global pandemic. Given the importance of T cells during host defense, strategies for TB vaccines largely aim at generating memory T cells rather than neutralizing antibodies. Most subunit vaccines incorporate immunodominant Mtb antigens, which elicit large T cell responses [6]. Several immunodominant antigens have been identified in the murine TB model, including Ag85a, Ag85b, CFP-10, ESAT-6 and TB10.4 [7]. T cell responses to these antigens are also frequently detectable in Mtb-infected people, and these highly prevalent responses represent the basis for TB immunodiagnostic tests [8]. By incorporating these immunodominant antigens into vaccines, the expectation is that antigen-specific T cells will contain the infection before Mtb can establish a niche and evade host immunity [6].

T cell recognition of Mtb*-*infected macrophages is fundamental to containment of TB infection. Srivastava et al elegantly showed this by using mixed bone marrow (BM) chimeric mice made from wild type (WT) and major histocompatibility complex class II (MHC class II) deficient BM [9]. Following infection, polyclonal CD4**^+^** T cells suppressed Mtb growth more efficiently in MHC class II-expressing cells than in MHC class II-deficient cells. This data convincingly argues that cognate recognition (i.e., T cell receptor (TCR) mediated recognition) of infected cells by polyclonal CD4**^+^** T cells limits bacterial growth. However, whether this protection comes from T cells recognizing immunodominant or subdominant antigens remains unknown. In fact, even though many presume that Mtb-infected cells present immunodominant antigens, the data validating this assumption is surprisingly inconsistent. While there is consensus that Mtb-infected cells present ESAT-6, the data concerning Ag85b presentation is more complicated [10-13]. Ag85b_241-256_ elicits a CD4**^+^** T cell response early after infection, but Mtb reduces Ag85b production within three weeks after in vivo infection [12]. Thus, while Ag85b_241-256_-specific CD4**^+^** T cells can recognize dendritic cells (DC) from infected mice 14 days post infection [14], there is little recognition of Mtb-infected cells by Ag85b_241-256_-specific CD4**^+^** T cells in vivo by day 21 [12]. Furthermore, Mtb has other mechanisms to evade T cell recognition, including dysregulating MHC class II expression and inhibiting antigen presentation by stimulating antigen export by the infected antigen presenting cells (APCs) [1, 12, 13, 15]. Whether the immunodominant antigens recognized by CD8**^+^** T cells are presented by Mtb-infected macrophages remains unknown. Here, we investigated cognate T cell recognition of Mtb-infected macrophages by CD8**^+^** T cells specific to the immunodominant antigen TB10.4.

TB10.4 (EsxH) is an ESAT-6-like protein secreted by the ESX-3 type VII secretion system, important in iron and zinc acquisition, and essential for Mtb growth in vitro and in vivo [16, 17]. Following Mtb infection, TB10.4 is a target of CD4**^+^** and CD8**^+^** T cell responses in humans and mice [18-22]. In Mtb-infected mice, TB10.4 elicits immunodominant responses in both BALB/c and C57BL/6 mice, and 30-50% of lung CD8^+^ T cells are specific to single epitopes (S1 Fig) [18, 19]. Whether these TB10.4-specific CD8^+^ T cells can mediate protection is unclear. Adoptive transfer of TB10.4-specific CD8^+^ T cells into Mtb-infected, immunocompromised mice reduces the bacterial burden and promotes host survival [19]. However, despite eliciting large numbers of TB10.4-specific CD8^+^ T cells, a vaccine incorporating the H-2 K^b^-restricted epitope, TB10.4_4-11_, fails to protect mice from Mtb infection [23]. We hypothesize that the inability of TB10.4-specific CD8^+^ T cells to mediate protection is due to inefficient recognition of Mtb-infected macrophages.

We used primary CD4**^+^** and CD8**^+^** T cells lines to investigate the recognition of Mtb-infected macrophages by T cells specific to Ag85b, ESAT-6, or TB10.4. Ag85b- and ESAT-6-specific CD4**^+^** T cells recognized Mtb-infected macrophages, but under the same conditions, TB10-specific CD8^+^ T cells did not recognize infected macrophages or inhibit bacterial growth. This was true even upon examination of numerous conditions and permutations including length of infection, duration of T cell and macrophage co-culture, and multiplicity of infection. TB10.4-specific CD8^+^ T cells did recognize macrophages infected with recombinant *Listeria monocytogenes* expressing TB10.4, but only if the bacilli could escape into the cytosol. However, overexpressing TB10.4 in Mtb did not confer recognition. Importantly, macrophages pulsed with irradiated bacteria efficiently cross-presented TB10.4 to CD8**^+^** T cells, suggesting that live Mtb actively inhibited presentation. Interestingly, polyclonal CD8**^+^** T cells specific for Mtb antigens other than TB10.4 recognized Mtb-infected macrophages in a MHC class I-restricted manner. Thus, while TB10.4-specific CD8**^+^** T cells do not recognize Mtb-infected macrophages, there exist other CD8**^+^** T cells that recognize subdominant antigens presented by Mtb-infected cells. Based on these data, we propose that TB10.4 is a decoy antigen: it elicits a massive and persistent CD8**^+^** T cell response, which cannot recognize Mtb-infected macrophages. Such a decoy antigen may distract the CD8**^+^** response from focusing on subdominant antigens presented by infected cells, leading to evasion from host immunity.

## Results

### TB10.4-specific CD8^+^ and Ag85b-specific CD4+ T cell lines sensitively recognize their cognate antigens

To study T cell recognition of Mtb-infected macrophages, we established antigen-specific T cell lines, which unlike T cell hybridomas, facilitate the study of T cell function as well as recognition. The TB10.4_4-11_-specific CD8**^+^** T cell line, referred hereafter as TB10Rg3, has a distinct TCR cloned originally from TB10.4_4-11_-tetramer^+^ CD8^+^ T cells isolated from infected mice and expressed in retrogenic mice [19]. The Ag85b240-254-specific CD4^+^ T cell line, referred hereafter as P25 cells, was derived from P25 TCR transgenic mice [24]. To confirm their antigen-specificity, we co-cultured the P25 or TB10Rg3 T cells with thioglycolate-elicited peritoneal macrophages (TGPMs) pulsed with or without their cognate peptides and then measured their expression of CD69 and Nur77. While both CD69 and Nur77 are T cell activation markers, increases in Nur77 expression indicate TCR-mediated activation more specifically [25, 26]. After co-culture with TGPMs pulsed with Ag85b_241-256_ peptide, Nur77 expression by P25 cells peaked after 2 hours (Fig 1a, b), while CD69 expression continued to increase (Fig 1c, d). TB10Rg3 T cells exhibited similar Nur77 and CD69 expression patterns after their co-culture with TGPMs pulsed with the TB10.4_4-11_ peptide (IMYNYPAM) but not with a control peptide (IMANAPAM) (Fig 1e-h). Since the increase in Nur77 expression was transient, we next tested whether CD69 and IFNγ could be useful markers of antigen recognition for longer experiments. During 72 hours of co-culture with peptide-pulsed TGPMs, P25 and TB10Rg3 T cells continued to express CD69 and secreted IFNγ in a peptide dose-dependent manner (Fig 1i-l). These experiments show that P25 and TB10Rg3 T cells can recognize their cognate antigens presented by TGPMs, both in short-term and long-term co-culture assays.

**Figure 1.**
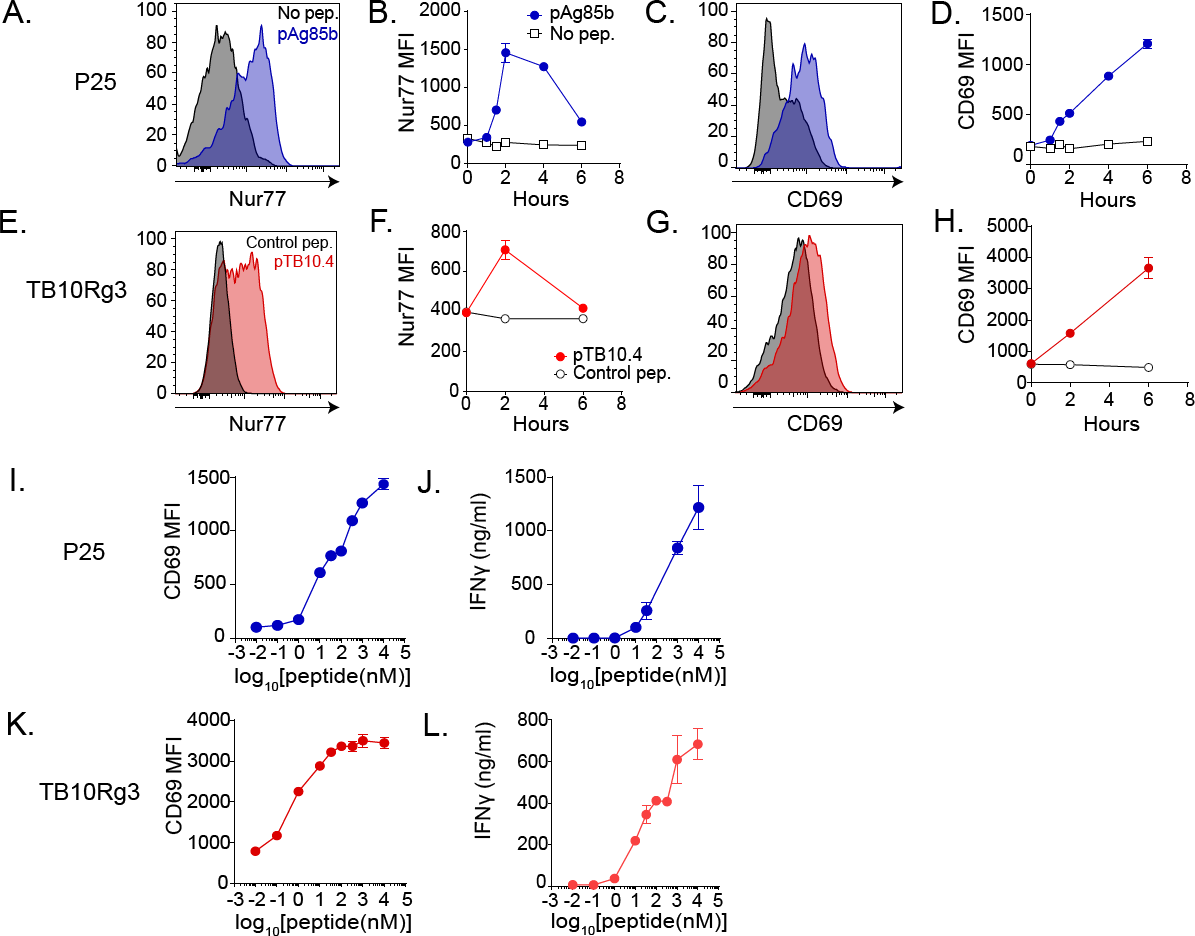
TB10.4-specific CD8^+^ (TB10Rg3) and Ag85b-specific CD4^+^ (P25) T cells both recognize their cognate peptides. (a) Representative histogram of Nur77 expression in P25 T cells after 2 hours of co-culture with macrophages and (b) time course of Nur77 MFI in P25 T cells. (c) Representative histogram of CD69 in P25 T cells after 2 hours of co-culture with macrophages and (d) time course of CD69 MFI in P25 T cells. (e) Representative histogram of Nur77 in TB10Rg3 T cells after 2 hours of co-culture and (f) time course of Nur77 MFI in TB10Rg3 T cells. (g) Representative histogram of CD69 in TB10Rg3 T cells at 2 hours of co-culture with macrophages and (h) time course of CD69 MFI in TB10Rg3 T cells. (i) CD69 MFI and (j) IFNγ production by P25 T cells after 72 hours of co-culture. (k) CD69 MFI and (l) IFNγ production by TB10Rg3 cells after 72 hours of co-culture. MFI, mean fluorescence intensity; mφ, macrophage.

### Ag85b-specific CD4^+^ T cells, but not TB10.4-specific CD8^+^ T cells, restrict intracellular bacterial replication

Given that a primary function of T cells during Mtb infection is to restrict bacterial growth, we determined whether these T cell lines could limit intracellular mycobacterial growth *in vitro*. We infected TGPMs with H37Rv, a virulent Mtb strain that expresses both TB10.4 and Ag85b in vitro [21, 27]. To assess whether any bacterial growth inhibition observed was dependent on cognate recognition, we infected both MHC-matched (i.e., H-2^b^) and mismatched (i.e., H-2^k^) macrophages. T cells were added on day 1 post-infection, and the number of colony forming units (CFU) was assayed 96 hours later. In the absence of T cells, Mtb grew significantly (p<0.01) (Fig 2). P25 T cells significantly inhibited intracellular bacterial growth in H37Rv-infected TGPMs (p<0.0001). Addition of Ag85b peptide to the infected macrophages did not further enhance the ability of P25 T cells to inhibit bacterial growth, suggesting that their activation was maximal. As expected, P25 T cells only inhibited bacterial growth in MHC-matched macrophages, indicating that growth inhibition mediated by T cells required cognate recognition under these conditions.

**Figure 2.**
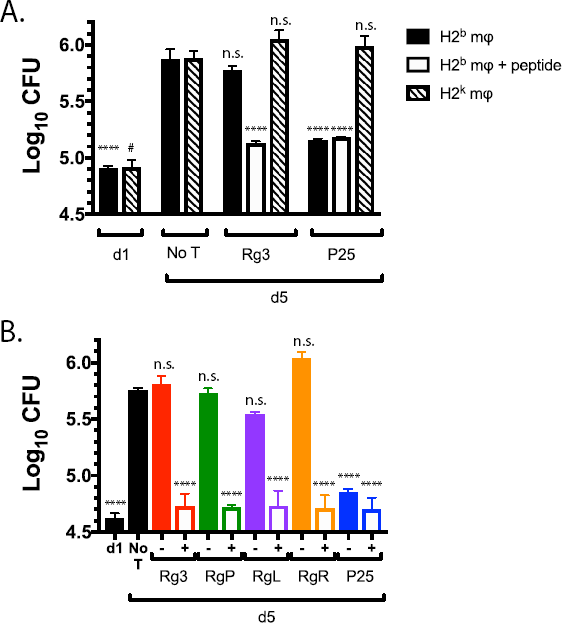
Ag85b-specific CD4^+^ T cells, but not TB10.4-specific CD8^+^ T cells, inhibit bacterial growth *in vitro*. (a) Matched (H-2^b^) or mismatched (H-2^k^) macrophages were infected with H37Rv for 2 hours, and then, one day post-infection, T cells were added. Separately, TB10.4 or Ag85b peptide was added to Mtb-infected (H-2^b^) macrophages, and then T cells were added. CFU were determined 4 days later. (b) C57BL/6 macrophages were infected with H37Rv as described above. The next day, TB10.4 or Ag85B peptide was added to selected wells. TB10Rg3, TB10RgP, TB10RgL, TB10RgR, or P25 T cells, were added, and CFUs were determined 4 days later. Results are representative of at least three experiments. Statistical testing was by one-way ANOVA, using the Dunnett posttest compared to d5. *, #, p<0.05; **, p<0.01; ***, p<0.005; ****, p<0.0001.

In contrast, TB10Rg3 T cells did not inhibit bacterial growth (Fig 2). We considered whether the inability of TB10Rg3 to inhibit bacterial growth was due to a lack of recognition of the infected macrophages or a defect in the T cells’ effector functions. When Mtb-infected TGPMs were pulsed with the TB10.44-11 peptide for one hour prior to adding the T cells, TB10Rg3 T cells significantly reduced bacterial growth (p<0.0001) (Fig 2a). Thus, under the same conditions where P25 T cells significantly suppressed intracellular Mtb growth in a MHC-restricted manner, TB10Rg3 T cells failed to inhibit bacterial growth.

We next considered whether the inability of TB10Rg3 to restrict intracellular bacterial growth was true for other TB10.4_4-11_-specific CD8^+^ T cells. To obtain TCRs representative of other clonally expanded TB10.4_4-11_-specific CD8^+^ T cells, TB10.4_4-11_-tetramer^+^ CD8^+^ T cells from Mtb-infected C57BL/6 mice were single cell sorted, and both the TCR CDR3α and CDR3β regions were sequenced (Fig S2). Three TCRs representative of clonally expanded T cells (TB10RgL, TB10RgR, and TB10RgQ) were cloned (Fig S2). In addition, a fourth TCR (TB10RgP), not previously identified by NexGen sequencing, was also cloned. All these TCRs responded specifically to the TB10.4_4-11_ epitope and expressed TCRs distinct from TB10Rg3 (Fig S3). We generated T cell lines from these retrogenic mice, as described for TB10Rg3. TB10RgP, TB10RgL, and TB10RgR did not inhibit bacterial growth (Fig 2b). However, if the Mtb-infected macrophages were pulsed with the TB10.44-11 peptide before T cell addition, then TB10RgP, TB10RgL, and TB10RgR T cells all inhibited bacterial growth significantly (Fig 2b). Since TB10.4_4-11_-specific CD8^+^ T cells only inhibited bacterial growth when their cognate peptide was added to Mtb-infected macrophages, we conclude that, although they express the effector function required to restrict intracellular bacterial growth, these TB10.4_4-11_-specific CD8^+^ T cells do not recognize Mtb-infected macrophages.

### Ag85b-specific CD4^+^ T cells, but not TB10.4-specific CD8^+^ T cells, recognize Mtb-infected macrophages

To further investigate TB10Rg3 and P25 T cells recognition of Mtb-infected cells, we next investigated the kinetics of Mtb antigen presentation. After Mtb infection, TGPMs were cultured for various lengths of time before adding the T cells. To assay antigen presentation, we added the T cells for two hours and then measured Nur77 and CD69 (see Fig 1 for kinetics). When added immediately after infection (i.e., day 0), P25 T cells recognized Mtb-infected macrophages based on the induction of Nur77 and CD69 (Fig 3a, b). Under these conditions, there was no increase in Nur77 or CD69 expression by TB10Rg3 T cells (Fig 3c, d). We next chose later time points, which might allow Mtb to adapt to the intracellular environment and potentially let the TB10.4 antigen accumulate. TB10Rg3 T cells were added to infected macrophages on days 1, 3, or 5 post-infection. Again, we did not observe any increase in Nur77 or CD69 expression (Fig 3e, f). As a control for T cell health and function, we co-cultured TB10.4_4-11_-peptide-pulsed-, uninfected-macrophages with the TB10Rg3 T cells and observed significant increases in their Nur77 and CD69 expression (Fig 3).

**Figure 3.**
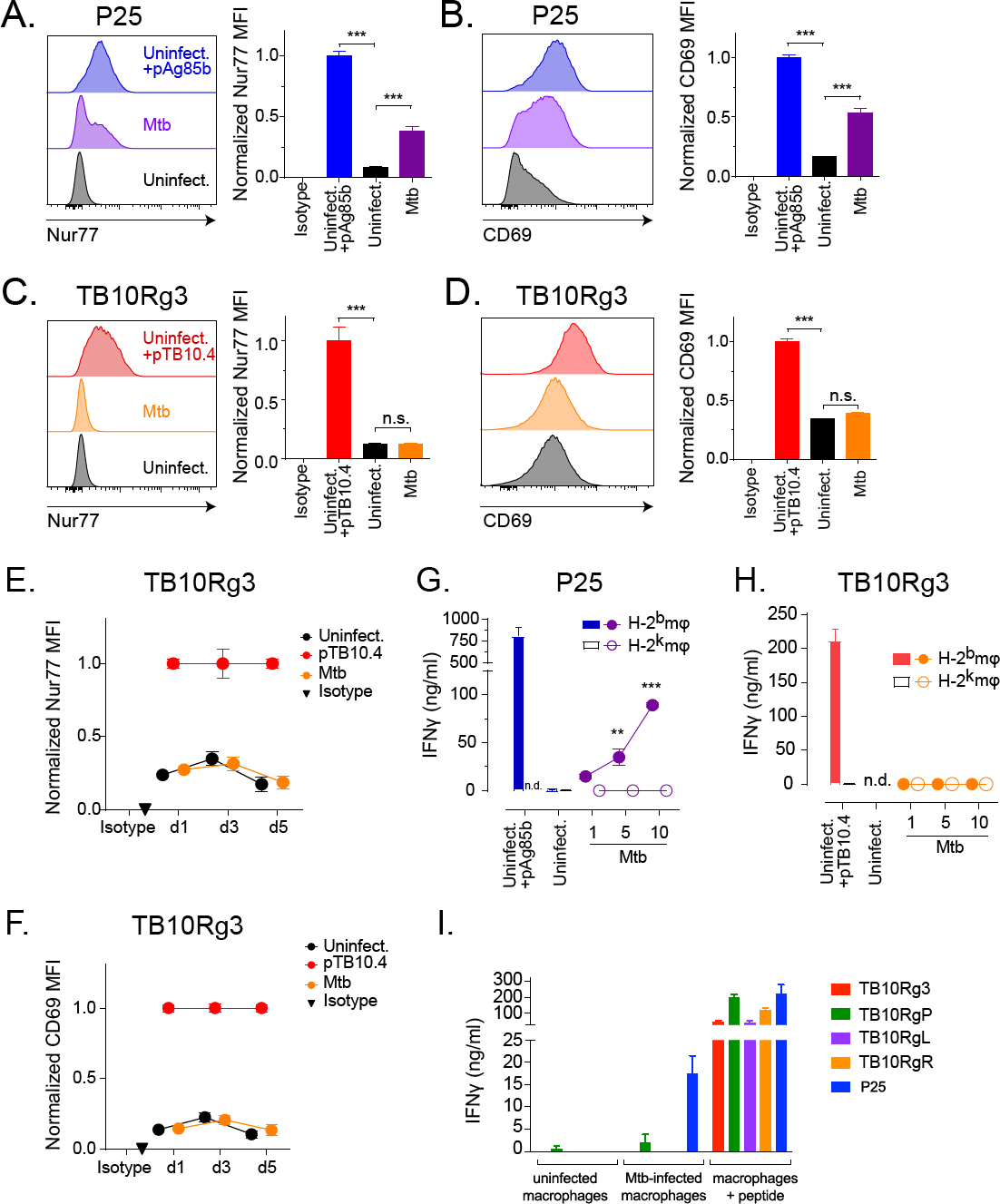
Ag85b-specific CD4^+^ T cells, but not TB10.4-specific CD8^+^ T cells, recognize Mtb-infected macrophages. (a-d) T cells were co-cultured with peptide-pulsed, Mtb-infected, or uninfected macrophages for 2 hours. (a) Representative histogram of Nur77 expression in P25 T cells and the normalized Nur77 MFI. (b) Representative histogram of CD69 expression in P25 T cells and the normalized CD69 MFI. (c) Representative histogram of Nur77 in TB10Rg3 T cells and the normalized Nur77 MFI. (d) Representative histogram of CD69 expression in TB10Rg3 T cells and the normalized CD69 MFI. (e-f) TB10Rg3 T cells were co-cultured with uninfected, peptide-pulsed, or Mtb-infected macrophages for 2 hours on d1, d3, and d5 post infection. Normalized expression of (e) Nur77 or (f) CD69 by TB10Rg3 T cells. P25 (g) or TB10Rg3 (h) T cells were co-cultured with uninfected, peptide-pulsed, or Mtb-infected macrophages, and IFNγ production was measured after 72 hours. (i) TB10Rg3, TB10RgP, TB10RgL, TB10RgR, or P25 T cells were cultured with uninfected, Mtb-infected, or peptide-pulsed macrophages. After 3 days, IFNγ in the supernatants was measured. Figures are representative of at least 5 (a-d, TB10Rg3), 2 (a-d, P25), 3 (e-h), or 2 (i) experiments. Statistical analysis was done by one-way ANOVA with Dunnett posttest (a-d) or two-way ANOVA with Sidak posttest (g-h). *, p<0.05; **, p<0.01; and ***, p<0.005.

Despite assessing recognition on multiple days, we considered whether the short assay period (i.e. 2 hours) might not detect recognition of Mtb-infected macrophages by TB10Rg3 T cells, especially if presentation of TB10.4 is inefficient or asynchronous. Therefore, we used IFNγ production as a cumulative indicator of T cell activation during a 72-hour co-culture experiment. Since cytokine-driven activation (e.g., IL-12, IL-18) can stimulate IFNγ production by T cells independently of TCR signaling, we used MHC-matched (H-2^b^) or mismatched (H-2^k^) TGPM to assess cognate recognition [26, 28-30]. As the infectious dose (MOI, multiplicity of infection) increased, P25 T cells produced more IFNγ when co-cultured with MHC-matched, but not MHC-mismatched, Mtb-infected TGPMs (Fig 3g). In contrast, TB10Rg3 T cells did not produce IFNγ when co-cultured with Mtb-infected TGPMs (Fig 3h). As before, TB10Rg3 T cells produced IFNγ when co-cultured with uninfected macrophages pulsed with the TB10.4_4-11_ peptide (Fig 3h).

We next used the TB10.4_4-11_-specific CD8^+^ T cell lines TB10RgP, TB10RgL and TB10RgR to address whether TB10.4_4-11_-specific CD8^+^ T cells other than TB10Rg3 can recognize Mtb-infected macrophages. While the TB10RgP, TB10RgL, and TB10RgR CD8^+^ T cell lines produced IFNγ when cultured with uninfected macrophages pulsed with the TB10.4_4-11_ peptide, none produced IFNγ following a 72-hour co-culture with Mtb-infected macrophages (Fig 3i). These data show that, regardless of the time point of T cell addition or the length of co-culture, P25 T cells, but not TB10.4_4-11_-specific CD8^+^ T cells, recognized Mtb-infected macrophages, based on their increased Nur77 and CD69 expression as well as their IFNγ production.

### TB10.4-specific CD8^+^ T cells do not recognize lung cells from Mtb-infected mice

During *in vivo* infection, Mtb infects a variety of myeloid cells, and this diversity changes over the course of the infection [31-33]. We considered that lung myeloid cells from Mtb-infected mice are more physiologically relevant than TGPMs. Thus, we isolated MHC class II^+^ lung cells from Erdman-infected, RAG-1-deficient mice 4 weeks post-infection and tested their ability to present Mtb antigens to TB10Rg3 T cells. We used RAG-1-deficient mice because of the possibility that CD8^+^ T cells in the lungs of Mtb-infected, wild type mice may recognize and eliminate any lung cells presenting the TB10.4 antigen. Since Mtb downregulates Ag85b expression by 3 weeks post infection [11, 12], we used an ESAT-6-specific CD4^+^ T cell line derived from C7 transgenic mice, which we refer to as C7 T cells [10, 34]. The immunodominant antigen ESAT-6 retains high levels of expression throughout infection and elicits a dominant CD4^+^ T cell response in C57BL/6 mice [11]. Due to the difficulty in obtaining large numbers of MHC class II^+^ cells from uninfected, RAG-1-deficient mice, we used TGPMs from age-matched, RAG-1-deficient mice as a source of uninfected, inflammatory macrophages. We stained C7 or TB10Rg3 T cells with 5uM of the proliferation dye eFluor450 (eBioscience) before co-culturing them with the lung myeloid cells. After 72 hours, we measured the T cell proliferation as a marker of T cell recognition. C7 T cells proliferated extensively when co-cultured with the infected lung myeloid cells but not when co-cultured with uninfected TGPMs (Fig 4a, b). In contrast, TB10Rg3 T cells did not proliferate when co-cultured with the lung myeloid cells (Fig 4c, d). To assess whether TB10Rg3 T cells could proliferate if TB10.4 was present, we pulsed the lung APCs with the TB10.44-11 peptide for 1 hour before adding the TB10Rg3 T cells. As predicted, TB10Rg3 T cells proliferated after 72 hours of co-culture with peptide-pulsed, lung myeloid cells (Fig 4c, d).

**Figure 4.**
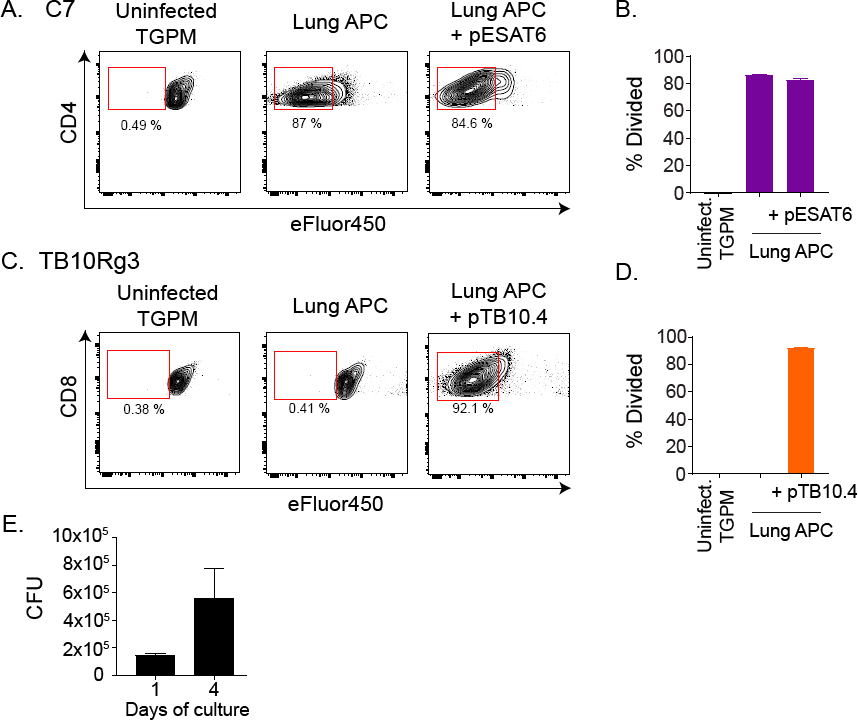
TB10Rg3 CD8^+^ T cells do not recognize lung APCs from infected mice. (a-d) T cell proliferation after coculture with lung APC from infected mice, with or without cognate peptide, or uninfected TGPM, based on eFluor450 fluorescence dilution after 72 hours. Representative flow plot (a) and quantification (b) of C7 T cell proliferation. Representative flow plot (c) and quantification (d) of TB10Rg3 T cell proliferation. (e) Bacterial burden in the lung APCs during *in vitro* culture over the course of the experiment in the absence of T cells. Representative of 4 (TB10Rg3) or 2 (C7) experiments.

We considered the possibility that Mtb in lung myeloid cells may not grow well in vitro, leading to altered antigen abundance that could affect T cell recognition. To address this possibility, we measured the bacterial burden in the lung myeloid cells. There was a 3-fold increase in the bacterial numbers between the beginning (d1) and the end (d4) of the experiment, indicating that the bacteria remained viable (Fig 4e). Together, these data indicate that, under the conditions in which C7 T cells recognized lung myeloid cells from Mtb-infected mice, TB10Rg3 T cells did not recognize these lung myeloid cells.

### TB10Rg3 CD8^+^ and P25 CD4^+^ T cells recognize macrophages infected with TB10.4-or Ag85b-expressing Listeria

We next investigated whether the location of the antigen might affect the presentation of TB10.4 since the MHC class I antigen presentation pathway primarily samples the cytosol, whereas Mtb is a classic phagosomal pathogen. TB10.4-specific CD8^+^ T cells are primed and expanded during Mtb infection, so the TB10.4 antigens must be cross-presented; however, whether Mtb-infected macrophages can competently cross-present mycobacterial antigens is unknown. We investigated these possibilities using DLLO or DActA mutant strains of *Listeria monocytogenes* engineered to express the full length TB10.4 protein, hereafter referred to as DLLO.TB10.4 or DActA.TB10.4, respectively. Both are attenuated strains: the DLLO.TB10.4 mutant cannot escape from the vacuole, while the DActA.TB10.4 mutant can escape from the vacuole but not from the cell. Hence, the TB10.4 protein made by the DLLO.TB10.4 strain will remain trapped in the phagosome, but the TB10.4 protein made by the DActA.TB10.4 strain will gain access to the cytosol.

TB10Rg3 T cells recognized DActA.TB10.4-infected TGPMs based on an increased frequency of Nur77-expressing cells (p<0.005) and the Nur77 MFI of all TB10Rg3 T cells (p<0.005) (Fig 5a-c). Bafilomycin, which inhibits vacuolar acidification and impairs the entry of the DActA.TB10.4 strain into the cytosol, diminished the frequency of Nur77-expressing cells (p<0.005) and Nur77 MFI (p<0.01) (Fig 5a, top, b, c). In contrast, TB10Rg3 T cells co-cultured with DLLO.TB10.4-infected TGPMs showed no increase in the frequency of Nur77-expressing cells or the Nur77 MFI (Fig 5a bottom, d, e). If recombinant listeriolysin (rLLO), the protein missing from the DLLO.TB10.4 strain, was added to the infected macrophages, an increase in the frequency of Nur77-expressing TB10Rg3 T cells (p<0.01) and the Nur77 MFI (p<0.01) became apparent. We also determined whether P25 T cells recognized Ag85b-expressing *Listeria monocytogenes* using the recombinant Listeria strains DActA.Ag85b and DLLO.Ag85b. Based on the propensity of MHC class II to present extracellular and vacuolar antigens, P25 cells recognized TGPMs infected with either DActA.Ag85b or DLLO.Ag85b, based on an increase in the frequency of Nur77-expressing T cells and Nur77 MFI (p<0.005) (Fig 5g, h). These results show that 1) TGPMs can efficiently process the full length TB10.4 protein and present the TB10.4_4-11_ epitope via MHC class I; 2) this process is more efficient when the bacteria are in the cytosol; and 3) TB10Rg3 T cells can efficiently recognize TB10.4_4-11_ presented during a live infection.

**Figure 5.**
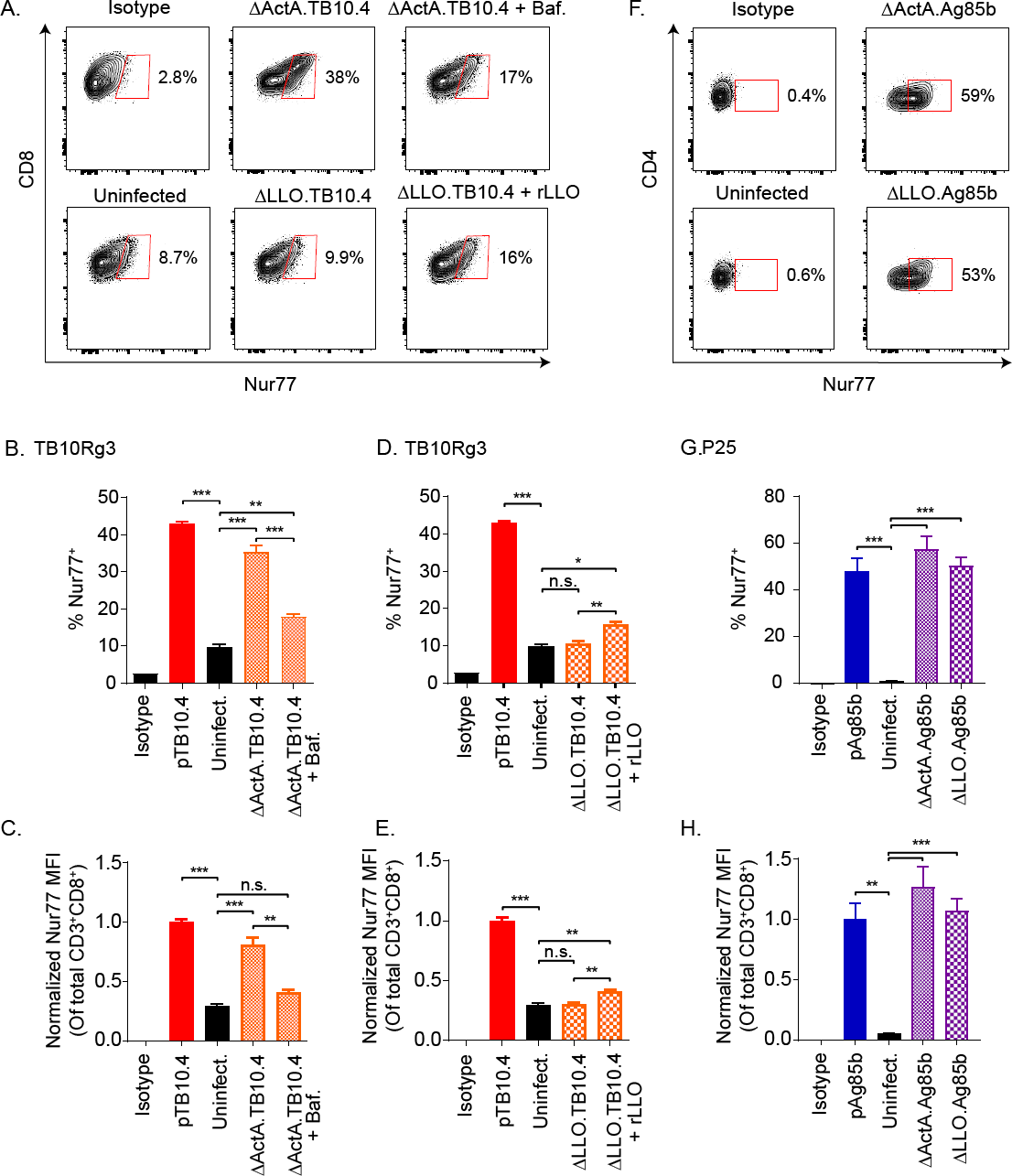
TB10Rg3 and P25 T cells can recognize macrophages infected with *Listeria monocytogenes* expressing TB10.4 and Ag85b proteins, respectively. (a) Representative flow plots showing Nur77 induction by TB10Rg3 T cells after co-culture with macrophages infected with DActA.TB10 (top row) or DLLO.TB10 (bottom row) Listeria. (b) Analysis of the frequency of Nur77-expressing TB10Rg3 T cells (b, d), or normalized MFI (c, e), after co-culture with DActA.TB10 (b, c) or DLLO.TB10 (d, e) infected macrophages. (f) Representative flow plots showing Nur77 induction by P25 population after co-culture with macrophages infected with DActA.TB10 (top row) or DLLO.TB10 (bottom row) Listeria. (g) Analysis of the frequency of Nur77-expressing P25 T cells and (h) normalized MFI of P25 T cells. Representative of at least two experiments. Statistical testing by one-way ANOVA with Dunnett posttest. *, p<0.05; **, p<0.01; and ***, p<0.005.

### TB10.4-specific CD8^+^ T cells do not recognize Mtb overexpressing TB10.4

We considered several additional possibilities as to why the TB10Rg3 T cells did not recognize Mtb-infected macrophages. Antigen abundance can affect T cell recognition, so we next tested whether increasing the level of TB10.4 protein expression might enhance TB10Rg3 T cell recognition of Mtb-infected macrophages. Since Mtb secretes esxH (TB10.4) together with esxG as a heterodimer [35], we developed a recombinant strain of H37Rv (esxGH-OE.Mtb), which overexpresses both esxG and esxH under the control of a tet^ON^ promoter. After tetracycline induction for 24 hours, the esxG and esxH mRNA expression increased multiple folds (Fig 6a). Prior to in vitro infection, we treated esxGH-OE.Mtb with or without tetracycline. The next day, TGPMs were infected with induced or uninduced esxGH-OE.Mtb. P25 T cells produced similar amounts of IFNγ when co-cultured with macrophages infected with either uninduced or induced esxGH-OE.Mtb, which was expected since Ag85b expression should not be altered (Fig 6b). Despite increasing the production of TB10.4 by Mtb, TB10Rg3 T cells still did not recognize Mtb-infected macrophages (Fig 6c). Although we cannot be certain that the induction of EsxGH leads to an increased amount of antigen delivered to the antigen processing pathway, this result suggests that antigen abundance is not limiting TB10.4-specific CD8^+^ T cell recognition of Mtb-infected macrophages.

**Figure 6.**
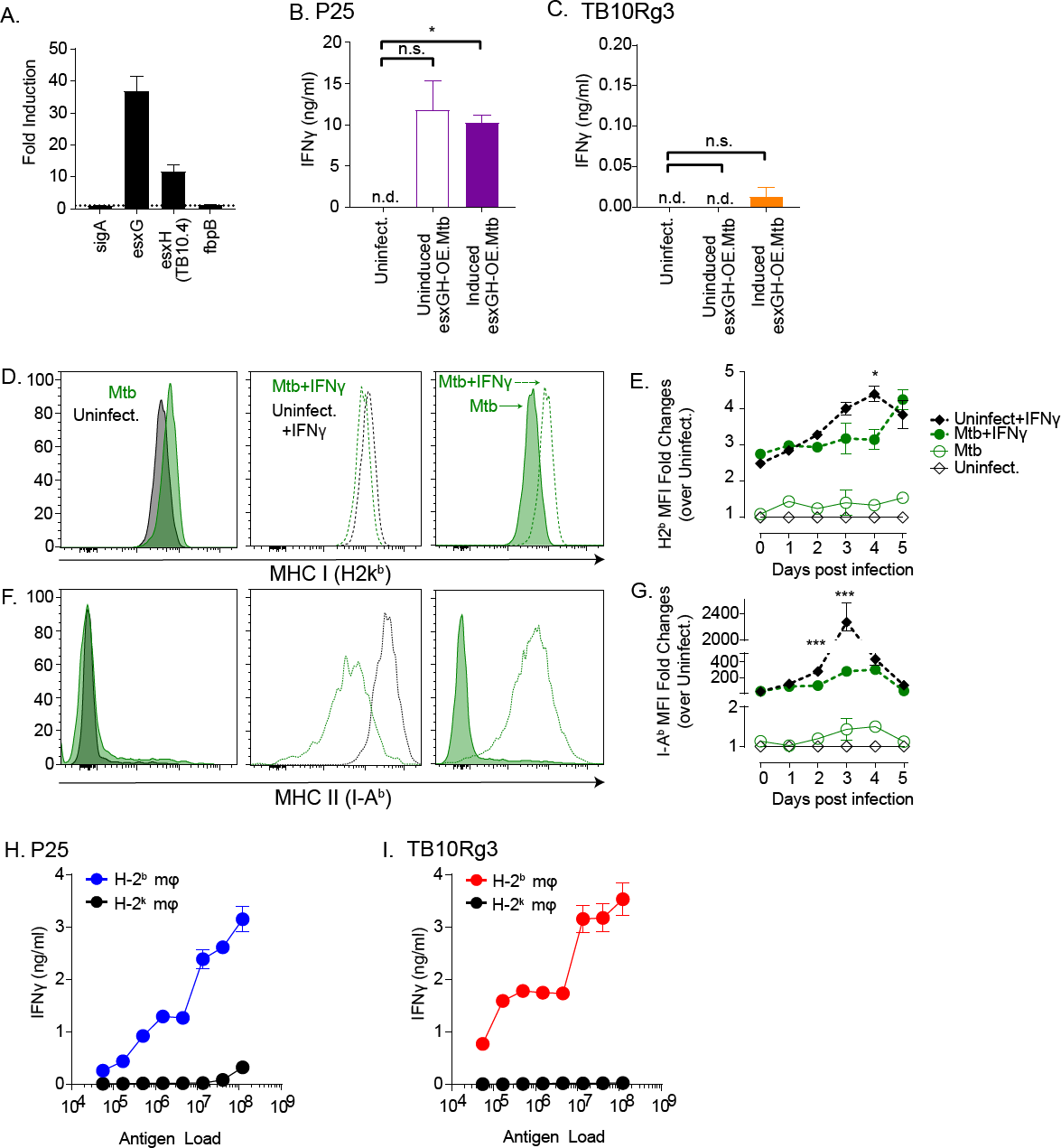
Probing potential mechanisms for lack of recognition by TB10Rg3 T cells. (a-c) EsxG (TB10.4) and its partner EsxH were overexpressed together in H37Rv to determine whether increasing TB10.4 abundance would lead to recognition of infected macrophages (esxGHOE.Mtb). (a) Tetracycline treatment of esxGH-OE.Mtb in broth culture induces esxG and esxH, but not fbpB and sigA, mRNA as measured by qPCR. Fold-induction was normalized to baseline (i.e., uninduced). IFNγ production by (b) P25 or (c) TB10Rg3 T cells after co-culture with macrophages infected with uninduced or induced esxGH-OE.Mtb. (d-g) MHC class I and II expression by Mtb infected-macrophages. Representative histograms (d, f) and (e, g) fold-change of MHC class I (d, e) or class II (f, g) expression on infected cells. (h) P25 and (i) TB10Rg3 production of IFNγ after co-culture with macrophages pulsed with titrated amounts of γ-irradiated (non-viable) H37Rv. Data is representative of 3 experiments. Statistical testing by one-way ANOVA with Dunnett posttest. *, p<0.05; **, p<0.01; and ***, p<0.005.

### Mtb infection does not significantly impair MHC class I and II expression by macrophages

We also investigated whether Mtb may inhibit MHC class I expression by infected TGPMs. Mtb and TLR2 agonists inhibit IFNγ-induced MHC class II expression by bone marrow derived macrophages, and the mycobacterial PPE38 protein can inhibit MHC class I expression in RAW264.7 macrophages and TGPMs infected with *Mycobacterium smegmatis* [36, 37]. Therefore, we asked whether Mtb impaired MHC class I expression in our in vitro infection system, especially since the TGPMs were not pre-activated with IFNγ prior to infection. We measured MHC class I and II expression by macrophages on each of the five days following infection. At baseline, uninfected TGPMs expressed high MHC class I, and Mtb infection did not alter MHC class I expression compared to the baseline (Fig 6d, e; solid lines). Although IFNγ pretreatment of macrophages led to an increase in MHC class I expression in uninfected TGPMs, infected TGPMs did not achieve the same peak levels (Fig 6d, e; dotted lines). As expected, the regulation of MHC class II was more sensitive to IFNγ. Uninfected TGPMs expressed low baseline levels of MHC class II (Fig 6f, g; solid lines). IFNγ pretreatment resulted in a >100-fold increase in MHC class II median fluorescence intensity (MFI) in the uninfected TGPMs, which peaked on day 3 with a >2000-fold increase over the baseline (Fig 6f, g; dotted lines). Mtb-infection alone did not significantly affect MHC class II expression, and consistent with previous studies, Mtb significantly impaired the induction of MHC class II by IFNγ pretreatment (Fig 6f, g). These data show that, in our in vitro infection model where the TGPMs were unstimulated, Mtb infection did not inhibit class I and II MHC expression. Importantly, the differences in MHC class I or class II expression by Mtb-infected macrophages cannot explain why P25 T cells, but not TB10Rg3 T cells, recognized infected macrophages.

### Macrophages cross-present antigens from non-viable Mtb to TB10.4-specific CD8^+^ T cells

Next, we hypothesized that Mtb may interfere with MHC class I presentation of mycobacterial antigens. Therefore, we tested the ability of the P25 and TB10Rg3 T cell lines to recognize TGPMs cultured with γ-irradiated, nonviable Mtb. Activation of pattern recognition receptors such as TLR2 and TLR4 by large amounts of dead bacteria might induce large amounts of IL-12 and IL-18, resulting in cytokine-driven T cell activation. Taking this concern into consideration, we used MHC-mismatched TGPMs as a control. We pulsed macrophages with a dose titration of γ-irradiated Mtb, then added TB10Rg3 or P25 T cells, and measured IFNγ secretion by the T cells after 72 hours. Both P25 and TB10Rg3 T cells produced high amounts of IFNγ when cultured with MHC-matched (i.e., H-2^b^) but not with MHC-mismatched (i.e., H-2^k^) TGPMs, and this response was dose dependent (Fig 6h, i). The ability of macrophages to process and present TB10.4 after phagocytosing γ-irradiated Mtb but not viable bacteria raises the possibility that live Mtb actively inhibit MHC class I presentation of TB10.4.

### Polyclonal, TB10.4_4-11_**-**tetramer negative CD8^+^ T cells from the lungs of Mtb-infected mice recognize infected macrophages

Along with the previous finding that TB10.4_4-11_-specific CD8^+^ T cells make up ∼40% of total lung CD8^+^ T cells during infection (S1 Figure) [19], our finding that TB10Rg3 T cells do not recognize Mtb-infected macrophages suggests that TB10.4 may be a decoy antigen. This raises the question whether the inability to recognize Mtb-infected macrophages is a general feature of the CD8^+^ T cell response to Mtb, or if this is a unique feature of TB10.4-specific CD8^+^T cells. Therefore, we determined whether polyclonal CD8^+^ T cells from the lungs of infected mice could recognize Mtb-infected macrophages. We carried out aerosol infection of C57BL/6 mice with Erdman, and, 6-8 weeks post infection, we purified polyclonal CD4^+^ or CD8^+^ T cells from their lungs and co-cultured them with Mtb-infected macrophages. After 72 hours of co-culture, polyclonal CD4^+^ T cells produced high amounts of IFNγ in a MHC-restricted manner (Fig 7a). Interestingly, polyclonal CD8^+^ T cells also produced IFNγ in a MHC-restricted manner when co-cultured with Mtb-infected macrophages (Fig 7b). These results indicate that other antigen-specific CD8^+^ T cells recognizing Mtb-infected macrophages do exist, and infected TGPMs can present Mtb antigens to CD8^+^ T cells. However, based on the high abundance of TB10.4-specific CD8^+^ T cells post infection (S1 Figure), the non-TB10.4-specific, Mtb-specific CD8^+^ T cells may be dwarfed by the dominant TB10.4-specific CD8^+^ T cells.

**Figure 7.**
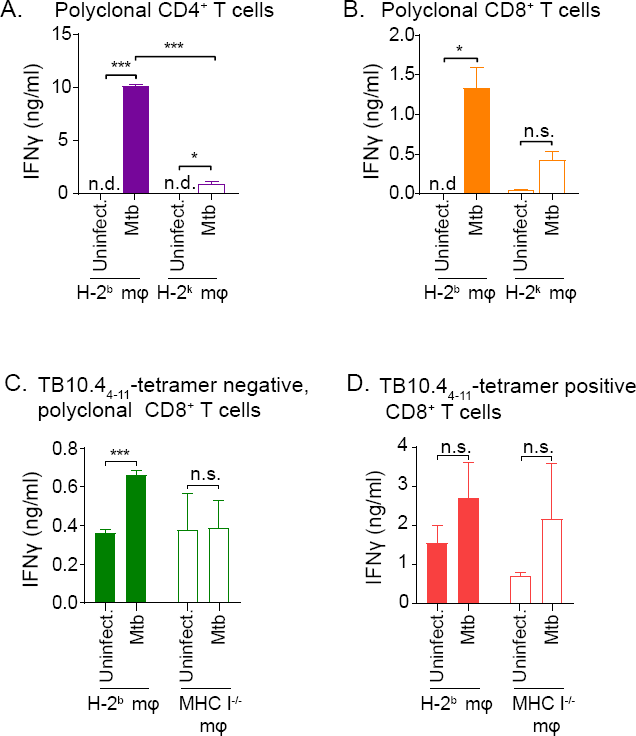
Polyclonal CD8^+^ T cells from the lungs of Mtb-infected mice recognize infected macrophages. IFNγ production by polyclonal CD4^+^ (a) or CD8^+^ (b) T cells after co-culture with either MHC-matched (H-2^b^) or MHC-mismatched (H-2^k^), Mtb-infected macrophages. IFNγ production by TB10.4_4-11_-tetramer-depleted (c) or tetramer-enriched (d) polyclonal CD8^+^T cells after co-culture with either MHC-matched (H-2^b^) or MHC-mismatched (H-2^k^), Mtb-infected macrophages. Data is representative of at least 2 experiments. Statistical testing by a two-tailed, unpaired Student’s T test. *, p<0.05; **, p<0.01; and ***, p<0.005.

To better assess whether the IFNγ production by polyclonal CD8^+^ T cells arose predominantly from non-TB10.4-specific CD8^+^ T cells, we used the TB10.4_4-11_-tetramer to separate TB10.4-specific and non-TB10.4-specific, polyclonal CD8^+^ T cells from the lungs of infected mice. After 72-hour co-culture with Mtb-infected macrophages, TB10.4_4-11_-tetramer negative CD8^+^ (non-TB10.4-specific CD8^+^) T cells produced significantly higher IFNγ compared to that of uninfected control (p<0.005), and the production was MHC class I restricted (Fig 7c). In contrast, TB10. 4_4-11_-specific CD8^+^ T cells produced IFNγ in a non-MHC-restricted manner during co-culture with both uninfected and Mtb-infected macrophages (Fig 7d). We cannot exclude the possibility that the tetramer isolation might have inadvertently activated the TB10.4_4-11_-specific CD8^+^ T cells. Nevertheless, these data show that polyclonal, TB10.4_4-11_-tetramer negative CD8^+^ T cells recognized Mtb-infected macrophages, supporting the notion of a subdominant T cell response that may be effective at detecting Mtb.

## Discussion

A complexity in defining T cell recognition is distinguishing cognate from non-cognate recognition. T cell IFNγ production, a common readout for recognition, can be stimulated by IL-12 and IL-18, two cytokines secreted by Mtb-infected cells [26, 28-30]. Even cognate recognition does not always signify recognition of infected cells. Uninfected macrophages and dendritic cells (DCs) can acquire exosomes, soluble proteins, apoptotic vesicles or necrotic debris containing non-viable bacilli or its antigens, and present these to T cells [13, 38-40]. This detour pathway allows T cells to be activated by uninfected DCs [38, 41]. Thus, T cell recognition of infected macrophages, which is central to our fundamental paradigm of TB pathogenesis, remains poorly defined.

Our study advances the understanding of T cell recognition of Mtb-infected cells. By focusing on TCR-mediated recognition, our data show that T cells specific to immunodominant antigens vary in their ability to recognize Mtb-infected macrophages. Despite being a persistent and dominant population of CD8^+^ T cells in the lungs of Mtb-infected mice, TB10.4_4-11_-specific CD8^+^ T cells do not recognize Mtb-infected macrophages. While we primarily used TGPMs, which have been used to model human macrophages [42, 43], we also showed that TB10.4-specific CD8^+^ T cells failed to recognize lung APCs from infected mice. Importantly, concurrent with our analysis of CD8^+^ T cells, we systematically assessed recognition of Mtb-infected macrophages by Ag85b-specific (i.e., P25) and ESAT-6-specific (i.e., C7) CD4^+^ T cells. Both recognized Mtb-infected macrophages and inhibited bacterial growth (here and [10]). Thus, under conditions that activated Mtb-specific CD4^+^ T cells, no activation of TB10.4-specific CD8^+^ T cells occurred. This finding has many implications, among which the most important is that not all Mtb-specific T cells recognize Mtb-infected macrophages.

These results led us to re-examine the evidence that CD8^+^ T cells recognize infected cells. In our evaluation of the literature, among the best evidence is: (1) direct ex vivo recognition of Mtb-infected macrophages and DC by CD4^+^ and CD8^+^ T cells [44-47]; (2) murine T cells’ cytolytic activity (CTL) of MTb-infected cells [48-50]; (3) human CD8^+^ T cells that recognize Mtb-infected DC [51-53]. However, these data have limitations. The murine studies never demonstrated cognate recognition, and the frequencies were lower than expected. The human studies only used DC and not macrophages and used a high MOI, raising concerns about death of infected cells and presentation of nonviable antigen. Nevertheless, these studies support the idea that CD8^+^ T cells recognize infected cells, but their frequency that recognize infected macrophages might be lower than we previously expected. Such a finding might explain why CD8^+^ T cells make a disproportionately small contribution to host defense, even though Mtb infection elicits a robust CD8^+^ T cell response.

We investigated several mechanisms that might explain why TB10.4-specific CD8^+^ T cells do not recognize infected macrophages. One possibility is the access of the TB10.4 antigen to the MHC class I processing pathway. Mtb can disrupt the phagosomal membrane and translocate into the cytosol [54], though this action often occurs later in infection and leads to necrosis of the macrophage [55]. We saw no evidence of recognition even at late time points such as days 4-5 post infection (Fig 3), when phagosomal disruption and bacterial translocation occurs [55]. The importance of antigen location became apparent during the Listeria infection experiments, where infected macrophages presented TB10.4_4-11_ only when the bacteria could enter the cytosol (i.e., DActA.TB10 but not DLLO.TB10). The Listeria experiments also provided an additional insight. Lindenstrom et al report that vaccination with TB10.4 (EsxH), which has a leucine at position 12 (i.e., IMYNYPAML), inefficiently generates TB10.4-specific CD8^+^ T cells [56]. However, vaccination with TB10.3 (EsxR), a related antigen that also contains the same epitope followed by a methionine (i.e., IMYNYPAMM), elicits TB10.4-specific CD8^+^ T cells. Based on that finding, they conclude there is a processing defect that prevents the generation of the TB10.44-11 epitope from the TB10.4 protein. However, they also find that TB10.4-specific CD8^+^ T cells elicited by TB10.3 vaccination recognize splenocytes pulsed with the rTB10.4 proteins, showing that the full length TB10.4 protein can be processed and presented. These data indicate that the lack of vaccine-elicited TB10.4-specific CD8^+^ T cells is due to a problem with priming after vaccination instead of an inability to process the IMYNYPAM epitope. Moreover, our data using TB10.4-expressing Listeria show that TGPMs can process the full length TB10.4 protein and present the TB10.44-11 epitope. Therefore, we conclude that amino acid sequence of TB10.4 does not hinder its processing. The Listeria experiments also show the potential importance of antigen location and raise the possibility that sequestration of the TB10.4 antigen in the phagosome renders it inaccessible to the MHC class I presentation pathway.

Low antigen abundance could also explain the lack of recognition. We have previously argued that there is limited amount of TB10.4 antigen presentation in the lungs of infected mice, leading to extreme bias in the TCR repertoire of the TB10.4-specific CD8^+^ T cell response and defects in the memory-recall response in vivo [19, 23]. To test the possibility of low antigen abundance, we overexpressed EsxG and EsxH (TB10.4) together but did not see greater T cell recognition of Mtb-infected macrophages, suggesting that abundance might not be the issue.

Unexpectedly, macrophages pulsed with γ-irradiated Mtb were recognized by TB10.4-specific CD8^+^ T cells, raising the possibility that live Mtb actively inhibits MHC class I presentation of TB10.4. This is particularly interesting since the presentation of CFP10, another ESAT-6-like protein, by human DCs to CD8^+^ T cells requires viable Mtb; DCs given heat-killed bacteria do not present CFP10 to T cells [53]. While these data suggest that presentation requires active secretion of CFP10 [57, 58], the heat-killing process could have destroyed CFP10, or there might not have been sufficient amounts of CFP10 available in the non-viable bacteria. In combination with our data showing polyclonal CD8^+^T cells recognize Mtb infected macrophages, these data show that it is possible that certain antigens are presented by live Mtb while others are actively prevented from being sampled by MHC class I.

Ag85b is an immunodominant antigen with an epitope recognized by CD4^+^ T cells in C57BL/6 mice. In vivo data shows that Ag85b-specific CD4^+^ T cells can recognize Mtb-infected cells early during infection; however, recognition decreases after infection is established [12, 14, 15, 59, 60]. The inability of Ag85b-specific CD4^+^ T cells to efficiently recognize Mtb-infected bone-marrow derived macrophages (BMDMs) or bone-marrow derived dendritic cells (BMDCs) stems from a combination of reduced Ag85b expression by Mtb and because infected cells actively export Ag85b into the extracellular milieu [12, 13]. In our experiments, we found that P25 T cells recognized Mtb-infected macrophages and inhibited bacterial growth in a MHC-restricted manner. A difference between the studies is the duration of macrophage and T cell co-culture. Grace et al examined 16-24 hours and found a lack of recognition, whereas our assays focused on 72-96 hours and detected recognition. Moreover, it is unknown whether Mtb-infected cells still exported antigens after the initial 24 hours of infection. Furthermore, the exported Ag85b could be taken up by infected cells during longer co-cultures, leading to their recognition by T cells. Finally, it is possible that cognate recognition of uninfected cells that present Ag85b could activate CD4^+^ T cells in a TCR-mediated manner, inducing IFNγ and indirectly controlling Mtb replication in macrophages. Nonetheless, under conditions that activate Mtb-specific CD4^+^ T cells, we could not observe activation of TB10.4-specific CD8^+^ T cells.

The TB10.44-11 epitope has been extensively used to characterize CD8^+^ T cell responses in the mouse model of TB, and TB10.4-specific CD8^+^ T cell responses have also been characterized in people with tuberculosis [19, 21, 23, 56, 61-63]. The finding that TB10.4-specific CD8^+^ T cells do not recognize infected macrophages was unexpected, particularly since TB10.4-specific CD8^+^ T cells persist in the lungs of infected mice and become more dominant with time [18, 19]. From these experiments, two questions warrant further investigation: 1) whether the CD8^+^ T cells specific to other epitopes of TB10.4 also inefficiently recognize infected macrophages, and 2) whether the species or the host genetic background influence recognition of infected cells.

In retrospect, our findings may partially explain why eliciting TB10.4-specific CD8^+^ T cells by vaccination fails to protect mice against Mtb infection [23, 56]. While vaccination with immunodominant antigens recognized by CD4^+^ T cells (e.g., Ag85b, ESAT-6) induce moderate protection [64, 65], we must consider the possibility that these antigens may not be the best stimulators of protective immunity. Ag85b-specific CD4^+^ T cells have variable efficacy, in large part due to its reduced expression by the bacterium as early as 3 weeks after infection [11, 12]. However, by their nature, the recruitment of memory T cell responses specific for immunodominant antigens is only incrementally faster than the primary T cell response [10, 23]. Thus, a crucial question for vaccine development is whether other Mtb antigens resemble TB10.4, in that they elicit T cell responses that fail to recognize infected macrophages. We did detect polyclonal CD8^+^ T cells that recognized Mtb infected macrophages, corroborating a previous study showing that polyclonal CD8^+^ T cells from infected mice can lyse Mtb-infected cells [48]. These data indicate that there are antigens presented by Mtb-infected cells, even if those antigens may be subdominant compared to TB10.4. Thus, future vaccine developments will benefit by identifying antigen targets based on their ability of being presented rather than only their immunogenicity.

Priming of TB10.4-specific CD8^+^ T cells occurs early after Mtb infection in the lung draining lymph node (LN) [23, 66]. Yet it is unknown whether priming of naïve T cells occurs via Mtb-infected DCs, or uninfected DCs that acquire antigen through uptake of apoptotic blebs containing Mtb proteins [38, 41], or by the transfer of antigen from cell to cell [40]. Priming by an uninfected cell can have detrimental consequences if the infected cell presents a different repertoire of Mtb antigens. Considering our findings, we propose that TB10.4 is a decoy antigen: TB10.4-specific CD8^+^ T cells expand in the LN during priming, accumulate in the lungs, but ineffectively recognize Mtb-infected macrophages. This raises the hypothesis that not all immune responses elicited by Mtb provide benefits to the host. Interestingly, Mtb genes encoding epitopes recognized by T cells are more highly conserved than other DNA elements, implying that T cell recognition of these Mtb epitopes may provide a survival advantage to the bacterium [67, 68]. For example, T cell dependent inflammation may benefit Mtb by promoting transmission. Even though TB10.4 is more variable than most other antigens, our results support these genetic data [67, 68]. Thus, Mtb focuses the CD8^+^ T cell response on the decoy antigen TB10.4 and distracts the immune response from other antigens that might be targets of protective immunity, successfully evading T cell immunity and enabling it to establish itself as persistent infection.

## Materials and Methods

### Ethics Statement

Studies involving animals were conducted following relevant guidelines and regulations, and the studies were approved by the Institutional Animal Care and Use Committee at the University of Massachusetts Medical School (Animal Welfare A3306-01), using the recommendations from the Guide for the Care and Use of Laboratory Animals of the National Institutes of Health and the Office of Laboratory Animal Welfare.

### Mice

C57BL/6J, Rag-1-deficient (B6.129S7-Rag1^tm1Mom^), B10 (C57BL/10J), B10.BR (B10.BRH2^k2^ H2-T18^a^/SgSnJJrep), P25 (C57BL/6-Tg(H-2K^b^-Tcrα,Tcrβ)P25Ktk/J) [24, 69], mice were obtained from Jackson Laboratories (Bar Harbor, ME). C57BL/6J and B10 mice were used for isolating MHC-matched TGPMs while B10.BR mice were used for isolating MHC-mismatched TGPMs. C57BL/6 K^b-/-^D^b-/-^ (MHC I^-/-^) mice were a generous gift from Dr. Kenneth Rock (University of Massachusetts Medical School, MA). C7 TCR transgenic mice were a generous gift from Dr. Eric Pamer (Memorial Sloan Kettering Cancer Center, NY)[34].

### Thioglycolate-elicited peritoneal macrophages

Thioglycolate-elicited peritoneal macrophages were obtained 4-5 days after intraperitoneal injection of mice with 3% thioglycolate solution, as described [28]. 1×10^5^ macrophages were plated per well. Macrophages were maintained in culture with RPMI 1640 media (Invitrogen Life Technologies, ThermoFisher, Waltham, MA) supplemented with 10 mM HEPES, 1 mM sodium pyruvate, 2 mM L-glutamine (all from Invitrogen Life Technologies) and 10% heat-inactivated fetal bovine serum (HyClone, GE Healthcare Life Sciences, Pittsburgh, PA), referred hereafter as supplemented complete media.

### Generation of CD8^+^ and CD4^+^ T cell lines

Retrogenic mice expressing TB10Rg3 TCR specific for the TB10.4_4-11_ epitope were generated as previously described [19]. The TB10Rg3 CD8^+^ T cells were isolated from these mice, stimulated *in vitro* with irradiated splenocytes pulsed with the peptide TB10.4_4-11_ in complete media containing IL-2. P25 or C7 CD4^+^ T cells were isolated from transgenic P25 or C7 mice, respectively [24, 34]. The P25 and C7 cells were stimulated *in vitro* with irradiated splenocytes pulsed with the Ag85b_241-256_ peptide or the ESAT-6_1-15_, respectively, in complete media containing IL-2 and anti-IL-4. After the initial stimulation, these T cells were split every two days for 3-4 divisions and rested for two to three weeks. After the initial stimulation, the cells were cultured in complete media containing IL-2 and IL-7.

### Peptides

The following synthetic peptide epitopes were used as antigens: TB10.4_4-11_ (IMYNYPAM); Ag85b_241-256_ (QDAYNAAGGHNAVFN); and ESAT-6_1-15_ (MTEQQWNFAGIEAAA). We also generated a negative control peptide predicted to not bind to H-2 K^b^: IMANAPAM. The peptides were obtained from New England Peptides (Gardner, MA).

As positive controls assessing the function of macrophages to present antigen, uninfected macrophages and, in certain experiments, infected macrophages were pulsed with the peptides of interest. We pulsed macrophages by incubating 10uM of the peptides of interest with the macrophages in supplemented complete RPMI 1640 media for 1 hour. After incubation, the cells were washed 3 to 5 times with fresh supplemented complete RPMI 1640 media. The cells were then resuspended in supplemented complete RPMI 1640 media for experiments.

### *In vitro* Mtb infection

H37Rv was grown as previously described [28]. Bacteria was grown to an OD_600_ of 0.6 – 1.0, washed in RPMI, opsonized with TB coat (RPMI 1640, 1% heat-inactivated FBS, 2% human serum, 0.05% Tween-80), washed again and filtered through a 5 micron filter to remove bacterial clumps. The bacteria were counted using a Petroff-Hausser chamber. Infection was performed as previously described [28]. The final multiplicity of infection (MOI), based on plating CFU, was 0.2-0.8 for all experiments. For CFU-based, bacterial growth inhibition assays, T cells were added at a ratio of 5 T cells to each macrophage. Four replicate wells were used for each condition. Cell cultures were lysed by adding 1/10^th^ volume of with 10% Triton X-100 in PBS (final concentration of 1%), and CFUs were determined by plating in serial dilutions of the lysates on Middlebrook 7H10 plates. CFUs were enumerated after culture for 21 days at 37°C and 5% CO_2_.

### *In vivo* aerosol Mtb infection and lung cell isolation

Aerosol infection of mice was done with the Erdman strain of Mtb using a Glas-Col aerosol-generation device. A bacterial aliquot was thawed, sonicated for 1 minute and then diluted in 0.9% NaCl-0.02% Tween-80 to 5 ml. The number of Mtb deposited in the lungs was determined for each experiment, by plating undiluted lung homogenate from a subset of the infected mice within 24 hours of infection. The inoculum varied between 37-120 CFU. For the ex vivo APC experiments, lung cells were isolated from Erdman-infected, RAG-1-deficient mice, 4-weeks post-infection, and the APCs were enriched by positive selection using anti-MHC class II^+^ microbeads (Miltenyi Biotec, Bergisch Gladbach, Germany) and the Miltenyi AutoMACS. On average, the isolated cells were 89% CD11c^+^ or CD11c^+^CD11b^+^. The APCs were counted on a hemocytometer and plated at 1×10^5^ per well in supplemented complete RPMI 1640 media.

For the ex vivo CD4^+^ and CD8^+^ T cell experiments, single cell suspensions were isolated from the lungs of infected C57BL/6 mice, 6 to 8 weeks post-infection, as described [10]. Polyclonal CD4^+^ and CD8^+^ T cells were enriched by positive selection using Mouse CD4^+^ and Mouse CD8^+^ T cell isolation kits, respectively (Miltenyi Biotec). After enrichment, average purity for polyclonal CD4^+^ and CD8^+^ T cells were 93% and 95%, respectively. For experiments investigating TB10.4_4-11_-tetramer positive cells and polyclonal, non-TB10.4-specific, CD8^+^ T cells, the following isolation was done. Single cell suspensions from the lungs of infected mice were incubated with APC-conjugated, TB10.4-_4-11_-loaded, H-2^b^ tetramers from the National Institute of Health Tetramer Core Facility (Emory University Vaccine Center, Atlanta, GA). Tetramer positive CD8^+^ T cells were then selected via the AutoMACS separator by anti-APC microbeads (Miltenyi Biotec). Average purity of TB10.4_4-11_-tetramer positive, CD8^+^ T cells was 85%, with 1.4% contaminating CD4^+^T cells. The tetramer negative population was subsequently washed and then enriched for polyclonal CD8^+^ T cells via Mouse CD8^+^ T cell isolation kit (Miltenyi Biotec). Average purity of polyclonal, non-TB10.4_4-11_-tetramer positive, CD8^+^ T cells was 75% with 0.8% contaminating CD4^+^T cells and 13% contaminating TB10.4_4-11_-tetramer positive CD8^+^ T cells. The T cells were counted using a hemocytometer and resuspended in supplemented complete RPMI 1640 media before being used in experiments.

### Listeria infections

The recombinant Listeria strains have been previously described [23]. For in vitro infections, they were grown to an OD_600_ of 0.6-1.0 in BHI media (Sigma Aldrich) with 10 ug/ml chloramphenicol (Sigma Aldrich) at 30°C. Macrophages were infected with the Listeria strains using a MOI 50, for 45 minutes. Extracellular bacteria were eliminated by adding 60 ug/ml gentamicin (Sigma Aldrich) for 20 minutes. Bacterial burden was assessed by lysing the infected macrophages with 1% TritonX-100 in PBS, and plating serial dilutions of the lysate on BHI agarose supplemented with 10ug/ml chloramphenicol (Sigma Aldrich, St. Louis, MO). Recombinant listeriolysin (Prospec, East Brunswick, NJ) was added in some experiments at 2 ug/ml for 30 minutes, and any excess was washed away. Bafilomycin (InvivoGen, San Diego, CA) was added in some experiments at 5 uM for 30 minute, before being washed away.

### Generation of TB10.4-overexpressing Mtb strains

pJR1103 was cleaved with EcoRI-HF and SalI-HF [70]. mCherry preceded by the groEL2 promoter from H37Rv was inserted by HiFi Assembly. The resulting plasmid was cleaved with NdeI and NotI-HF. The esxGH gene from H37Rv, along with 12 upstream nucleotides, was inserted by HiFi Assembly following the plasmid-borne tetracycline-inducible promoter. All enzymes used above were purchased from New England Biolabs. The resulting plasmid (pGB6) was electroporated into Mtb H37Rv and integrated at the L5 site. RNA was purified from induced and uninduced cultures using TRIzol (ThermoFisher) and chloroform extraction, followed by purification on Zymo columns. cDNA was produced with Superscript IV (ThermoFisher), and quantitative PCR was performed using the iTaq SYBR Green Supermix (Bio-Rad, Hercules, CA) on an Applied Biosystems Viia 7 thermocycler.

### Irradiated H37Rv

The following reagent was obtained through BEI Resources, NIAID, NIH: *Mycobacterium tuberculosis*, Strain H37Rv, Gamma-Irradiated Whole Cells, NR-14819. The irradiated H37Rv was gently sonicated using a cup-horn sonicator at a low power to disperse bacterial clumps while limiting bacterial lysis. The number of bacteria were approximated by measuring the turbidity at OD_600_, and correlating it with live H37Rv (OD_600_ = 1 is equivalent to 3.0×10^8^ CFU/ml). To pulse TGPMs, diluted, sonicated, γ-irradiated H37Rv were added to adherent macrophages for one hour before repeatedly washing the cultures to remove residual extracellular bacteria. Subsequently, TB10Rg3 or P25 T cells were added at a ratio of 1 T cell to 1 macrophages. After 72 hours, the amount of IFNγ in the supernatants was measured using Mouse IFNγ ELISA MAX kits (Biolegend, San Diego, CA).

### Flow Cytometry Analysis

The following cell surface antigens were detected by flow cytometry using the following antibodies: mouse CD4 (clone GK1.5), CD8 (clone 53-6.7), CD3ε (clone 145-2C11), CD69 (clone H1.2F3), I-A/I-E (clone M5/114.15.2), and H-2K^b^ (clone AF6-88.5) (all from Biolegend). BV421-and APC-conjugated, TB10.4_4-11_-loaded, H-2K^b^ tetramers were obtained from the National Institutes of Health Tetramer Core Facility (Emory University Vaccine Center, Atlanta, GA). Zombie Violet Fixable viability dye (Biolegend) or the Live/Dead Fixable Far Red Dead Cell stain (ThermoFisher) were used for distinguishing live from dead cells. To stain for the Nur77 transcription factor, the Nur77 monoclonal antibody (clone 12.14) was used in combination with the Foxp3 Transcription Factor Staining Buffer Set (both from ThermoFisher) by the manufacturer’s protocol. Live/dead viability staining and surface staining were done for 20 minutes at 4°C, and intracellular staining was done for 30 minutes at room temperature. Samples were then fixed with 1% paraformaldehyde/PBS for 1 hour before being analyzed by a MACSQuant flow cytometer (Miltenyi Biotec). FlowJo Software (Tree Star, Portland, OR) was used to analyze the collected data. Single lymphocytes were gated by forward scatter versus height and side scatter for size and granularity, and dead cells were excluded.

### Normalization and statistical analysis

To compare the cellular expression of Nur77 and CD69 expression levels between time points, the MFI values were normalized as follows: experimental values were divided by the difference between the isotype control MFI (minimum response) and the peptide control MFI (maximum response).

Each figure represents a minimum of 2 similar experiments, with 2 to 4 biological replicates in each experiment. Data are represented as mean ± standard error of the mean (SEM). For comparing two groups, a two-tailed, unpaired student’s t-test was used. For more than two groups, the data were analyzed using a one-way ANOVA. A p value < 0.05 was considered to be statistically significant. Analysis was performed using GraphPad Prism, Ver. 7 (GraphPad Software, La Jolla, CA).

## Supplemental Legends

**S1 Supplemental Figure.**
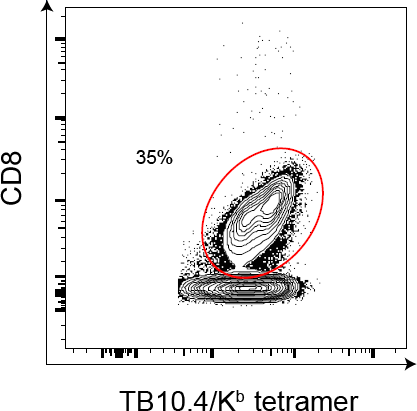
TB10.4_4-11_**-**tetramer positive CD8^+^ dominates the pulmonary CD8^+^ T cell response during Mtb infection in C57BL/6 mice. Representative flow plot showing the percent of TB10.4_4-11_-tetramer positive CD8^+^T cells among lung cells isolated from mice infected with Erdman via the aerosol route 6 weeks post-infection.

**S2 Supplemental Figure:**
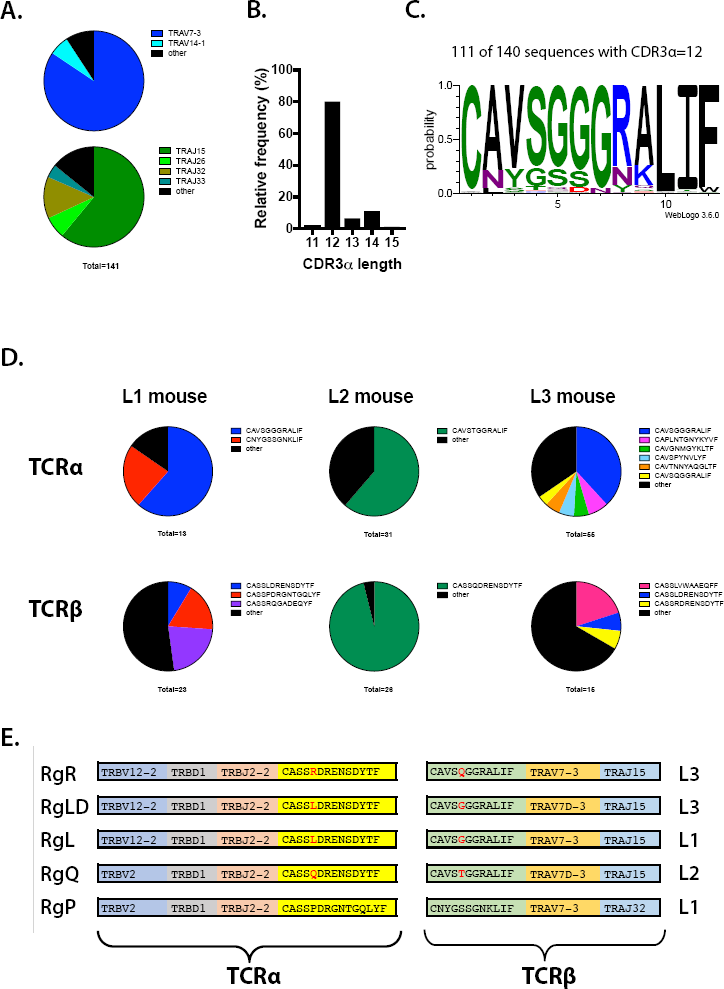
TCR analysis and cloning of TCRs expressed by TB10.4_4-11-_ specific CD8^+^ T cells from the lungs of Mtb-infected C57BL/6 mice. Mononuclear lung cells were obtained from three individual C57BL/6 mice, 9 weeks after low-dose aerosol infection with Mtb Erdman. Single TB10.4_4-11_-tetramer^+^ CD8^+^ T cells from each mouse were sorted into 96-well plates and the CDR3α and CDR3β sequences determined as described [19]. The analysis of a representative mouse is shown, for which 141 CDR3α sequences was determined. A clonal expansion of TB10.4_4-11_-tetramer^+^CD8^+^ T cells, as previously described [19], was suggested by the skewed distribution of TRAV and TRAJ families (a), which shows an extreme bias in the use of TRAV7 and TRAJ15 gene segments, as well as a dominant CDR3α amino acid (aa) length of 12 (b). (c) Analysis of all CDR3α aa sequence with a length of 12 (n=112) identify a consensus motif of CAVSGGGRALIF for TB10.4_4-11_-specific CD8^+^ T cells. Amplification of CDR3α and CDR3β sequences from the same well allowed pairing of TCRα and TCRβ for individual TB10.4_4-11_-specific CD8^+^ T cells. Three individual mice were analyzed in this manner (d). We identified an expanded CDR3β sequence containing the “xDRENSD” motif, the same motif that had been previously defined by NexGen sequencing [19]. Thus, mouse L1 had an expansion of CD8^+^ T cells with the CASSLDRENDYTF CDR3β sequence, mouse L2 was dominated by CD8^+^ T cells using the CDR3β sequence CASSQDRENDYTF, and mouse L3 expressed two major expansions, one encoding CASSLDRENDYTF and the other, CASSDDRENDYTF (d). Based on our ability to pair the CDR3α and CDR3β sequences, we detected an interesting reciprocal conservation. Namely, the ‘xDRENSD’ CDR3β motif was matched to a ‘SxGGRA’ CDR3α motif (e). Finally, we identified an expansion of a T cell clone in mouse L1, which expressed a novel sequence that we had not previously observed (i.e., CASSPDRGNTGQLYF) (d, e). Thus, with a high degree of confidence, we paired the CDR3α and CDR3β sequences belonging to 5 distinct TB10.4_4-11_-specific CD8^+^ T cell clones that had been expanded in lungs of Mtb-infected C57BL/6 mice. The TCRα and TCRβ cDNAs were reconstructed and cloned using standard methods, and retrogenic mice were subsequently produced [19, 71, 72].

**S3 Supplemental Figure.**
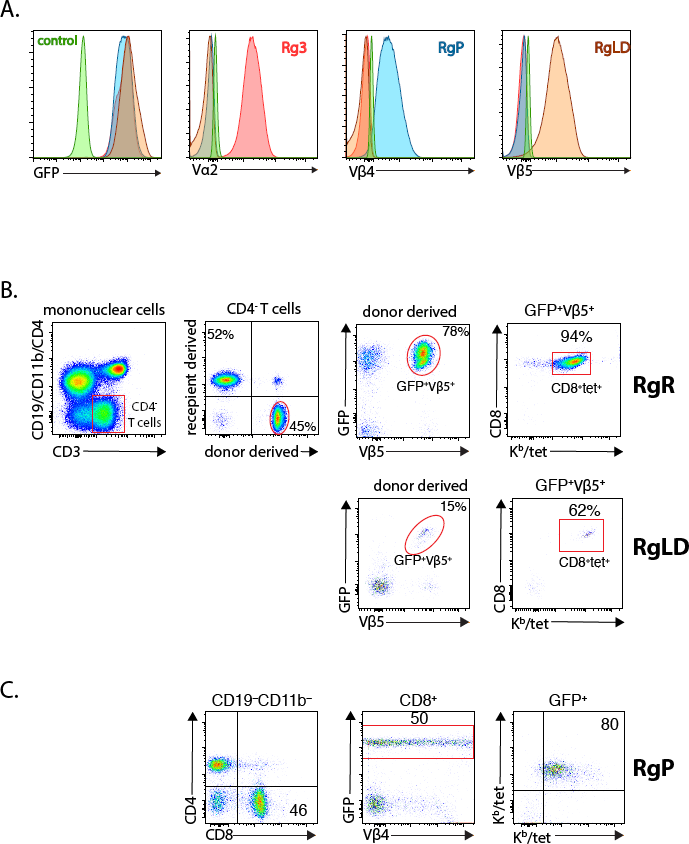
Reconstitution and expression of specific TCRs in C57BL/6 retrogenic mice. Retrogenic mice were produced as previously described [19]. Six weeks after retroviral transduction of bone marrow and reconstitution of congenically marked recipient mice, the expression of the recombinant TCR was determined in peripheral blood. (a) The BW58α^-^β^-^ cell line was transduced with different retroviral constructs. GFP^+^ cells were sorted three times, and mAbs specific for Vα or Vβ were used to confirm successful TCR expression and pairing of TB10RgP and TB10RgLD. The TB10Rg3 construct was included as internal control. (b) Representative flow cytometry plots showed gating strategy of donor-derived GFP^+^ specific Vβ^+^ TB10.4_4-11_-tetramer^+^ CD8^+^ TB10RgR and TB10RgLD mice. (c) Representative flow cytometry plots of splenic T cells from TB10RgP retrogenic mice demonstrating CD8^+^GFP^+^ T cells staining with the TB10.4_4-11_-tetramer.

## Acknowledgements

We would like to thank Dr. Kenneth Rock (University of Massachusetts Medical School, MA) for the generous gift of C57BL/6 K^b-/-^D^b-/-^ (MHC I^-/-^) mice. We thank Barry Kriegsman for critical reading of the manuscript. We also thank the Animal Medicine team at University of Massachusetts Medical School (Worcester, MA) for their technical assistance. This study was funded by grants from the National Institutes of Health (R01 AI106725, R01 AI123286).

